# Fungal seed pathogens of wild chili peppers possess multiple mechanisms to tolerate capsaicinoids

**DOI:** 10.1101/712711

**Authors:** Catharine A. Adams, Kolea Zimmerman, Kristi Fenstermacher, Mitchell G. Thompson, Will Skyrud, Scott Behie, Anne Pringle

**Author notes:** Corresponding author: Catharine Adams.

## Abstract

The wild chili pepper *Capsicum chacoense* produces the spicy defense compounds known as capsaicinoids, including capsaicin and dihydrocapsaicin, antagonistic to the growth of fungal pathogens. Compared to other microbes, fungi isolated from infected seeds of *C. chacoense* possess much higher tolerance to these spicy compounds, having their growth slowed, but not entirely inhibited. Previous research has shown capsaicinoids inhibit microbes by disrupting ATP production via the binding of NADH dehydrogenase in the Electron Transport Chain (ETC), throttling Oxidative Phosphorylation (OXPHOS). Capsaicinoids may also disrupt cell membranes. Here, we investigated capsaicinoid tolerance in fungal seed pathogens isolated from *C. chacoense*. We selected 16 fungal isolates from four *Ascomycete* genera (*Alternaria, Colletotrichum, Fusarium* and *Phomopsis)*. Using relative growth rate as a readout for tolerance, fungi were challenged with ETC inhibitors to infer if fungi possess alternative respiratory enzymes, and if effects on the ETC fully explained inhibition by capsaicinoids. In all isolates, we found evidence for at least one alternative NADH dehydrogenase. In many isolates we also found evidence for an alternative oxidase. These data suggest wild plant pathogens may be a rich source of alternative respiratory enzymes. We further demonstrate these fungal isolates are capable of the breakdown of capsaicinoids. Lastly, we determine the OXPHOS theory weakly explains the primary mechanism by which dihydrocapsaicin slows fungal growth, but not capsaicin. Our findings suggest capsaicinoids likely disrupt membranes in addition to energy poisoning, with implications for microbiology and human health.

**Importance:** Plants make chemical compounds to protect themselves. For example, chili peppers produce the spicy compound capsaicin to inhibit animal feeding and pathogen damage. In humans, capsaicin binds to a membrane channel protein, creating the sensation of heat, while in microbes, capsaicin limits energy production by binding respiratory enzymes. However, some data suggest capsaicin also disrupts membranes. Here we studied fungal pathogens (*Alternaria*, *Colletotrichum*, *Fusarium*, and *Phomopsis*) isolated from a wild chili pepper, *Capsicum chacoense*. By measuring growth rate in the presence of antibiotics with known respiratory targets, we infer wild plant pathogens may be rich with alternative respiratory enzymes. A zone of clearance around the colonies, as well as LCMS data, further indicate these fungi can break down capsaicin. Lastly, the total inhibitory effect of capsaicin was not fully explained by its effect on respiratory enzymes. Our findings lend credence to studies proposing capsaicin may disrupt cell membranes, with implications for microbiology as well as human health.

## Introduction

Successful seed dispersal is critical to plant fitness (Alejandro Estrada and Theodore H. Fleming 1986). Many plants coat seeds in fleshy fruits to attract dispersers, but the high density of sugar and nutrients in fruits also attracts fungal and bacterial pathogens (Kolb et al. 2007). To deter pathogens, plants make a diversity of secondary metabolites, including phenolics and polyphenols, terpenoids, essential oils, alkaloids, lectins, polypeptides and more (Cowan 1999). Many plants produce these secondary metabolites directly in the fruit to surround the seeds, such as the various amides in *Piper* fruits *(Whitehead and Bowers 2014)*, caffeine in coffee fruits (Almeida et al. 2006), and capsaicinoids in chili peppers.

*Capsicum* is an anthropologically and economically important plant genus, largely because of its secondary metabolite production (Qin et al. 2014). The genus is unique in its production of capsaicinoids, the alkaloids that give chili peppers their spicy pungency (Naves et al. 2019). Early American people began eating chilies over 6000 years ago, and cultivated the plants around 3000 BCE (Perry et al. 2007). After European discovery of the Americas, pungent chili peppers rapidly became a staple dish in many cuisines around the world (Andrews 1993), at least partially due to the antimicrobial properties of capsaicinoids such as capsaicin (Billing and Sherman 1998).

Plant secondary metabolites, such as capsaicinoids, can protect against novel pathogens (Cipollini and Levey 1997), but microbes that commonly associate with plants can evolve resistance to defensive compounds (Wink 1988), the result being a co-evolutionary arms race between plants and pathogens (Thompson and Burdon 1992; Burdon and Thrall 1999).

*Capsicum* plants make capsaicinoids in the placenta of the fruit, which surrounds the seeds (Aza-González et al. 2011; Reyes-Escogido et al. 2011) and reduces the growth of fungal seed pathogens of a wild chili species, *C. chacoense* (Tewksbury et al. 2008). Though fungal growth is slowed in the presence of capsaicinoids, these fungi exhibit an extreme tolerance to capsaicinoids relative to other tested microbes (Jones et al. 1997; Xing et al. 2006; Chatterjee et al. 2010). However, the exact mechanism of capsaicin tolerance in these wild fungi is unknown.

Two lines of research have examined the antimicrobial effects of the primary capsaicinoid, capsaicin. The first body of research has assessed the ability of capsaicin to disrupt cell membranes, and has shown capsaicin can affect the structure and phase organization of model membranes (Aranda et al. 1995). Capsaicin also affects voltage-dependent sodium channels and alters bilayer elasticity in artificial membranes (Lundbaek et al. 2005), but the effects of capsaicin on microbial cell membranes have not yet, to our knowledge, been tested *in vivo*.

A separate, larger body of work examines the ability of capsaicin to inhibit energy production. In bacteria and fungi, capsaicin binds to Complex I of the Electron Transport Chain (ETC), thus inhibiting oxidative phosphorylation (OXPHOS) and subsequent energy (ATP) production (Shimomura et al. 1989; Yagi 1990) (Figure 1). *Saccharomyces cerevisiae* and many other yeasts are largely insensitive to capsaicin, presumably because they lack Complex I, the NADH dehydrogenase (Yagi 1990). Instead, yeasts possess alternative NADH dehydrogenases. These less efficient alternative respiratory enzymes lack the energy-coupling site to push protons into the mitochondrial intermembrane space, but can still transfer electrons from NADH to downstream complexes (Joseph-Horne et al. 2001) (Figure 1).

**Figure 1.**
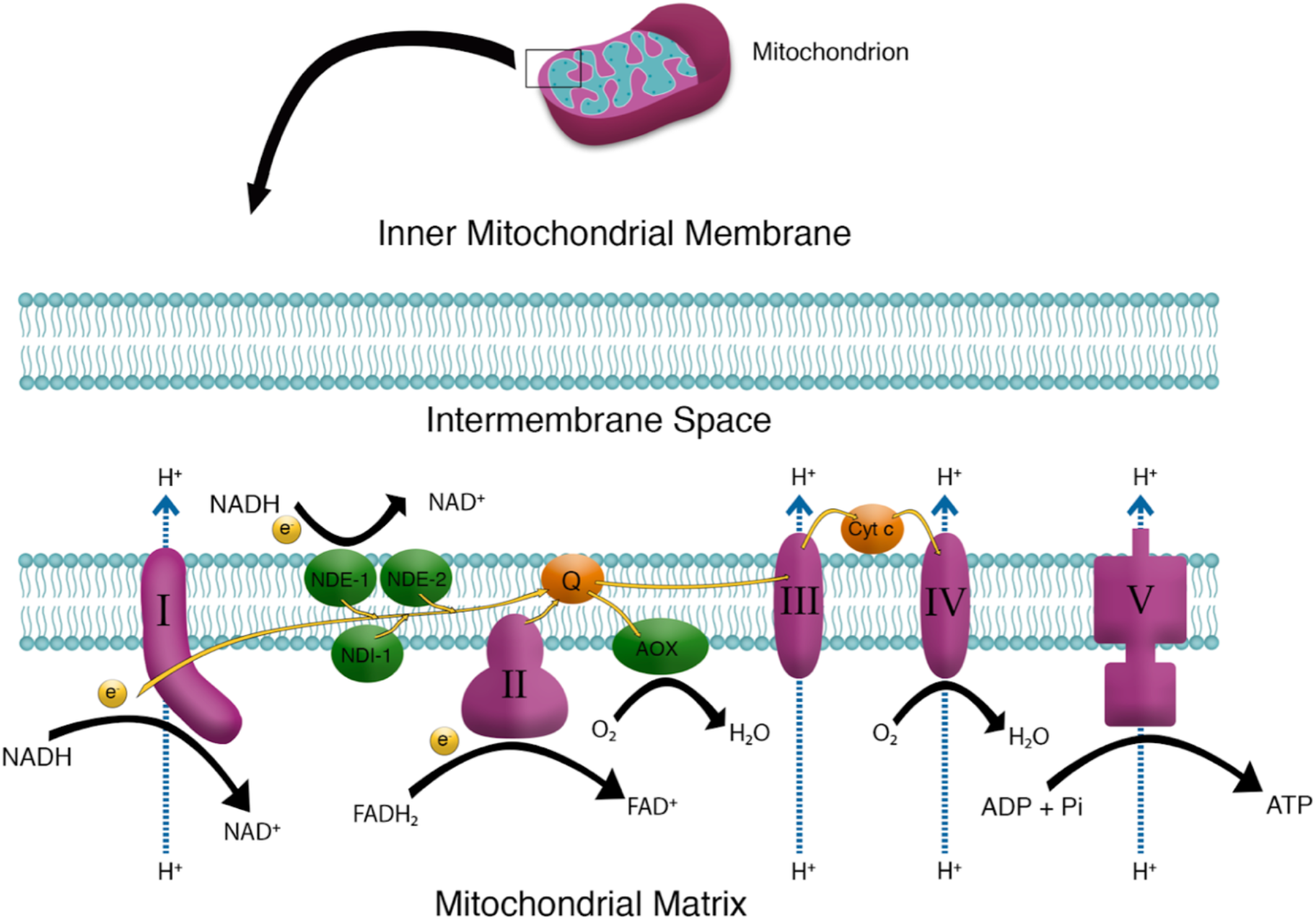
The standard and alternative respiratory enzymes of a fungal electron transport chain. Standard respiratory enzymes are shown in pink. The standard Complexes both accept electrons and pump protons to generate proton motive force. The entire standard electron transport chain includes Complex I, the NADH dehydrogenase; Complex II, the succinate dehydrogenase; membrane-embedded ubiquinone (Q); Complex III, the cytochrome bc_1_ complex; soluble cytochrome c (C); Complex IV, the cytochrome c oxidase; and Complex V, the ATP synthase. Alternative respiratory enzymes named from *Saccharomyces cerevisiae* are shown in green. Three alternative Complex I enzymes are known in fungi: Nde1 and Nde2, which embed in the external membrane; and Ndi1 in the internal membrane, facing the mitochondrial matrix. These alternative respiratory enzymes accept electrons from NADH but do not pump protons into the intermembrane space. The alternative oxidase (AOX) can directly transfer electrons from ubiquinol to oxygen, reducing oxygen to water, but bypasses electron transfer through Complexes III and IV, reducing overall proton motive force and subsequent ATP production, compared to the standard complexes.

Other microbes possess alternative NADH dehydrogenase enzymes, in addition to Complex I: such enzymes have been described in the malaria parasite *Plasmodium yoelii yoelii (Uyemura et al. 2004)*, in the causal agent of corn smut *Ustilago maydis*, and in the industrially important filamentous fungus *Aspergillus niger* (O’Donnell et al. 2011). Many fungi possess genes for alternative respiratory enzymes (Marcet-Houben et al. 2009), but the physiological function of these enzymes has only been tested in a few model fungi, such as the yeast *Yarrowia lipolytica* (Kerscher et al. 1999) and the filamentous fungus *Neurospora crassa (Carneiro et al. 2007)*.

Previous work with alternative respiratory enzymes has taken advantage of ETC inhibitors that block standard respiratory enzymes, but are rescued by alternative enzymes. These ETC inhibitors allow researchers to test for the presence of alternative enzymes in non-model organisms with relatively intractable genetic systems (Table 1). For example, the drug rotenone inhibits Complex I, the standard NADH dehydrogenase, but has no effect on alternative NADH dehydrogenases (Joseph-Horne et al. 2001; Juárez et al. 2004; Kerscher et al. 1999). Thus, fungal growth in the presence of this inhibitor indicates the presence of an alternative NADH dehydrogenase.

**Table 1.** ETC inhibitors used in this study, and their known targets within the mitochondrion.

| Inhibitor | Known target(s) in fungi | Concentration used in this study |
| --- | --- | --- |
| Antimycin | Complex III. Does **not** affect AOX. | 5 micromolar |
| Capsaicin | Complex I. Does **not** affect alternative Complex I enzymes. | 50 micromolar |
| Chloramphenicol | Inhibits mitochondrial translation, standard and alternative Complex I enzymes, and alternative oxidase. | 7.7638 millimolar[Office1] |
| Dihydrocapsaicin | Complex I. Does **not** affect alternative Complex I enzymes. | 50 micromolar |
| Flavone | Nde1, Nde2, and Ndi1. Does **not** target Complex I. | 0.5 millimolar |
| Oligomycin | Complex V. | 5 micromolar |
| Rotenone | Complex I. Does **not** affect alternative Complex I enzymes. | 5 micromolar |

In addition to alternative respiratory enzymes to bypass the effects of capsaicinoids, fungi that associate with chili peppers may possess degradative enzymes that can decrease the concentration of the capsaicinoids they encounter. An enzyme that degrades capsaicinoids has been found in several bacteria, but to our knowledge, only one fungus, *Aspergillus oryzae (Lee et al. 2015)*. The bacterial degradative capacities include an echinocandin acylase from *Actinoplanes utahensis* NRLL 12052 (Romano et al. 2011), *Bacillus* species isolated from Korean kimchi (Cho et al. 2014), and an acylase from *Streptomyces mobaraensis* (Koreishi et al. 2006).

In this study, we measured the growth of fungi isolated from infected seeds of the wild Bolivian chili pepper, *C. chacoense.* For these experiments, we selected fungi from four genera known to be pathogenic on *Capsicum* cultivars in agricultural systems: *Alternaria, Colletotrichum, Fusarium*, and *Phomopsis*, for a total of 16 isolates—four fungi from each genus. First, we measured fungal growth in the presence or absence of ETC inhibitors (Table 1) to infer whether these fungi possess alternative Complex 1 enzymes, and if these fungi possess alternative Complex 3 enzymes. Using a combination of agar plate based degradation studies and LC-MS, we evaluated whether fungal isolates could degrade capsaicin. Finally, we grew fungi on a fermentable (glucose) or a non-fermentable (glycerol) carbon source to investigate whether the inhibitory effect of capsaicinoids on respiratory enzymes wholly explained their effect.

## Materials and Methods

### Collection sites and fungal isolation

All fungi were isolated from fruits collected from previously established study sites in the Gran Chaco region, Bolivia (Tewksbury et al. 2008).

We isolated fungi from infected seeds by transferring one seed per fruit to agar plates that contained dilute potato dextrose media (PDA) with a 0.2% solution of the antibiotics tetracycline, ampicillin, and streptomycin. We incubated seeds in the dark at 25° C and checked them daily for fungal growth. Isolates were subcultured until they were axenic, transferred onto agar slants for storage, and maintained at 6° C until use.

We selected 16 total isolates that could be assigned to one of four different genera: *Fusarium (Sordariomycetes: Hypocreales; Phomopsis (Sordariomycetes: Diaporthales); Colletotrichum (Sordariomycetes: Glomeralleles);* and *Alternaria (Dothidiomycetes: Pleosporales)*. Within each fungal genus studied, we selected four fungal isolates that had the highest capsaicin tolerance when grown on false fruit media (data not shown). Aside from isolate Alt1, which was contaminated and could not be re-isolated, the isolates used in this study will be deposited in the Fungal Genetics Stock Center, and subcultures of all isolates in the collection can be supplied upon request.

### DNA extraction and sequencing

To determine the identity of fungal isolates, we grew all the fungi in our 200+ culture collection on Potato Dextrose agar prior to DNA extraction and PCR of the ITS1 region. We amplified the ITS1 5.8S ITS2 region of nuclear ribosomal DNA with 1 µl each of the 25µM ITS5 (5’ GGAAGTAAAAGTCGTAACAAGG 3’) and Nl4 (5’ GGTCCGTGTTTCAAGACGG 3’). PCR protocol details are described elsewhere (Machnicki 2013). We submitted samples for automatic sequencing in both directions to the Penn State Genomic Core Facility, University Park, PA on an ABI 3739XL (Applied Biosystems, Foster City, CA). We edited sequence data and assembled consensus sequences using Sequencher (Gene Codes Corporation, Ann Arbor, Michigan, USA).

We estimated taxonomic placement at the genus level for each isolate by BLAST searching the NCBI GenBank nucleotide database (http://www.ncbi.nlm.nih.gov/BLAST/). To assign a genus, we used the highest GenBank sequence similarity scores available, and assigned a genus name if the sequence similarity score was 90% or higher and the Expect-value (E-value) was 0.0.

### Tested compounds

We used the following drugs, purchased from the listed manufacturers: the complex I inhibitors capsaicin (8-methyl-*N*-vanillyl-6-nonenamide, Sigma-Aldrich), dihydrocapsaicin (N-(4-Hydroxy-3-methoxybenzyl)-8-methylnonanamide, Sigma-Aldrich), and rotenone (2R,6aS,12aS)-1,2,6,6a,12,12a-hexahydro-2-isopropenyl-8,9-dimethoxychromeno[3,4-b] furo(2,3-h)chromen-6-one, Sigma-Aldrich); complex II inhibitor antimycin ([(2R,3S,6S,7R,8R)-3-[(3-formamido-2-hydroxybenzoyl)amino]-8-hexyl-2,6-dimethyl-4,9-dioxo-1,5-dioxonan-7-yl] 3-methylbutanoate, Sigma-Aldrich); complex V inhibitor oligomycin ((1S,2’R,4E,5’R,6R,6’S,7S,8R,10S,11S,12R,14S,15R,16S,18E,20E,22S,25R,28R, 29S)-22-ethyl-3’,4’,5’,6’-tetrahydro-7,11,14,15-tetrahydroxy-6’-[(1Z)-2-hydroxy-1-propen-1-yl]-5’,6,8,10,12,14,16,28,29-nonamethyl-spiro[2,26-dioxabicyclo[23.3.1]nonacosa-4,18,20-triene-27,2’-[2H]pyran]-3,9,13-trione, Sigma-Aldrich); mitochondrial protein synthesis inhibitor chloramphenicol (2,2-dichloro-N-[1,3-dihydroxy-1-(4-nitrophenyl)propan-2-yl]acetamide, CalBioChem); and alternative NADH dehydrogenase inhibitor flavone (2-Phenyl-4*H*-1-benzopyran-4-one, 2-Phenylchromone, Sigma-Aldrich) (Martins et al. 2011). The targets of the drugs are listed in Table 1.

### Tests of antifungal activities

#### a) Carbon sources

To pinpoint the effects of each drug on OXPHOS via the ETC, we used two different carbon sources. Glucose can be fermented and used in glycolysis, while glycerol, which cannot be fermented, is only used in OXPHOS (Sun et al. 2013) (Sun et al. 2013).

#### b) Preparation of the ETC inhibitors

All the test compounds (sans chloramphenicol; see below) were dissolved in dimethylsulfoxide (DMSO), sterilized by filtration using a 0.2 µmeter VWR Sterile Syringe Filter (28145-501), and kept as stock solutions.

The targets of all the drugs, and the concentrations of the drugs used, are listed in Table 1. To determine a relevant concentration of capsaicin and dihydrocapsaicin to use in experiments, we referred to previous work with HPLC assays of *Capsicum chacoense* wild fruits (Haak et al. 2012). Pungent fruits possess capsaicinoids in amounts of roughly 0.5 mg/g dry weight. This concentration moderately inhibits fungal growth without causing complete inhibition (Tewksbury et al. 2008). Thus to mimic this ecologically relevant concentration, we added capsaicinoids at a concentration of 0.25 mg/ml, or 50 micromolar (Table I).

Antimycin, oligomycin, and rotenone were each added to the media at a concentration of 5 micromolar. Flavone was added at a concentration of 0.5 millimolar, as in (Uyemura et al. 2004).

A subset of fungi showed growth on glycerol at every concentration of chloramphenicol tested, so we selected the highest concentration in which chloramphenicol is soluble in water (7.7368 millimolar). To avoid confounding effects of high volumes of DMSO solvent, unsterilized chloramphenicol powder was added directly to the autoclaved, liquid media. We observed no contamination.

#### c) Fungal growth assays

For all treatments, we dispensed 5 ml of each growth media into 60 mm petri dishes. We grew fungi on minimal media with 1.5% agar, and 0.0555 molarity of carbon from either glucose or glycerol. To standardize the size of the initial inoculum, we used sterile 5 mm Pasteur pipettes to excise and transfer plugs onto each plate. On each carbon type, each isolate was plated on a set of experimental drug plates as well as solvent and plain-media control plates. Solvent control plates contained 2 ml of the drug solvent DMSO per 750 ml of media, and were used to separate the effects of drugs from possible toxic effects of the drug solvent. To quantify any possible effects of DMSO, we also made negative controls with no DMSO added (plain-media). Plain-media controls lacked all drugs and solvents.

Precultures were grown on plates with 20 ml of potato dextrose agar (PDA). We replicated each drug treatment three times for each of the 16 isolates, with solvent and carbon controls, for a total of 832 plates. Plates were kept in the dark at 25° C. At 48 hours after inoculation, we measured two perpendicular colony diameters, and noted the time of measurement. We used the mean colony diameter for each isolate, and calculated colony diameter increase in units of mm/hr.

#### d) Confirmation of degradation with LC-MS

To confirm whether the observed zone of clearance on capsaicinoid plates (Supplementary Figure S1A) was truly indicative of capsaicin degradation, isolate Fus3 was selected for further investigation. Plates were made with either 100 µmolar capsaicin or 0 µmolar capsaicin, and inoculated with fungus (Fungal) or not (Control), in triplicate. After 10 days of growth, one 5 mm plug was excised from the edge of fungal growth (or equivalent) and transferred to a 2 ml eppendorf tube. Two volumes of ice-cold methanol were then added to the sample and stored at −20° C until analysis.

LC-UV-MS analysis was performed on an Agilent 6120 single quadrupole LC-MS equipped with an Agilent Eclipse Plus C18 column (4.6 × 100 mm). A linear gradient of 2-98% CH_3_CN with 0.1% formic acid (v/v) over 30 min in H_2_O with 0.1% formic acid (v/v) at a flow rate of 0.5 mL/min was used. Samples were monitored at 280 nm using a coupled Agilent 1260 infinity DAD. Extracted ion chromatograms were integrated and peak area was used to construct a standard curve using an authentic capsaicin standard. Concentrations of capsaicin within samples were interpolated from this curve.

LC-HRMS analysis was performed on an Agilent 6545 quadrupole time of flight LC-MS equipped with an Agilent Eclipse Plus C18 column (4.6 × 100 mm). A linear gradient of 2-98% CH_3_CN with 0.1% formic acid (v/v) over 30 min in H_2_O with 0.1% formic acid (v/v) at a flow rate of 0.5 mL/min was used. Samples were also monitored at 280 nm using a coupled Agilent 1290 infinity DAD.

### Data Analysis

We considered the effect of each drug when fungi were grown on either fermentable glucose or non-fermentable glycerol. On each carbon type, we calculated the percent inhibition of each drug by subtracting the average growth rate on solvent control plates from the average growth rate on treatment plates, and then dividing by the average growth rate on solvent control plates. For chloramphenicol, which was not dissolved in DMSO, we compared growth to media controls.

Statistical analyses were conducted using the R language version 3.0.2 (The R Core Team 2013). We calculated growth rate with the lubridate package (Grolemund and Wickham 2011), and plots were generated with the ggplot2 package (Wickham 2009). Propagation of error was used to construct the error bars in Figure 5. For correlation analyses we used linear models, and we used one-sided t-tests to compare between averages, except in the case of figure 6, for which we used a two-sided t-test.

## Results

### Fungal pathogens of wild chili peppers possess alternative respiratory enzymes

To infer the presence of an alternative Complex 1 enzyme, we tested fungal growth rate when inhibited by the Complex I inhibitor rotenone. When grown on non-fermentable glycerol, the fungal isolates showed only 4.83% inhibition by rotenone (Figure 2A). Conversely, flavone was 62.86% more effective on average than rotenone (p < 0.0005), with an average inhibition of 67.69% (Figure 2B).

**Figure 2.**
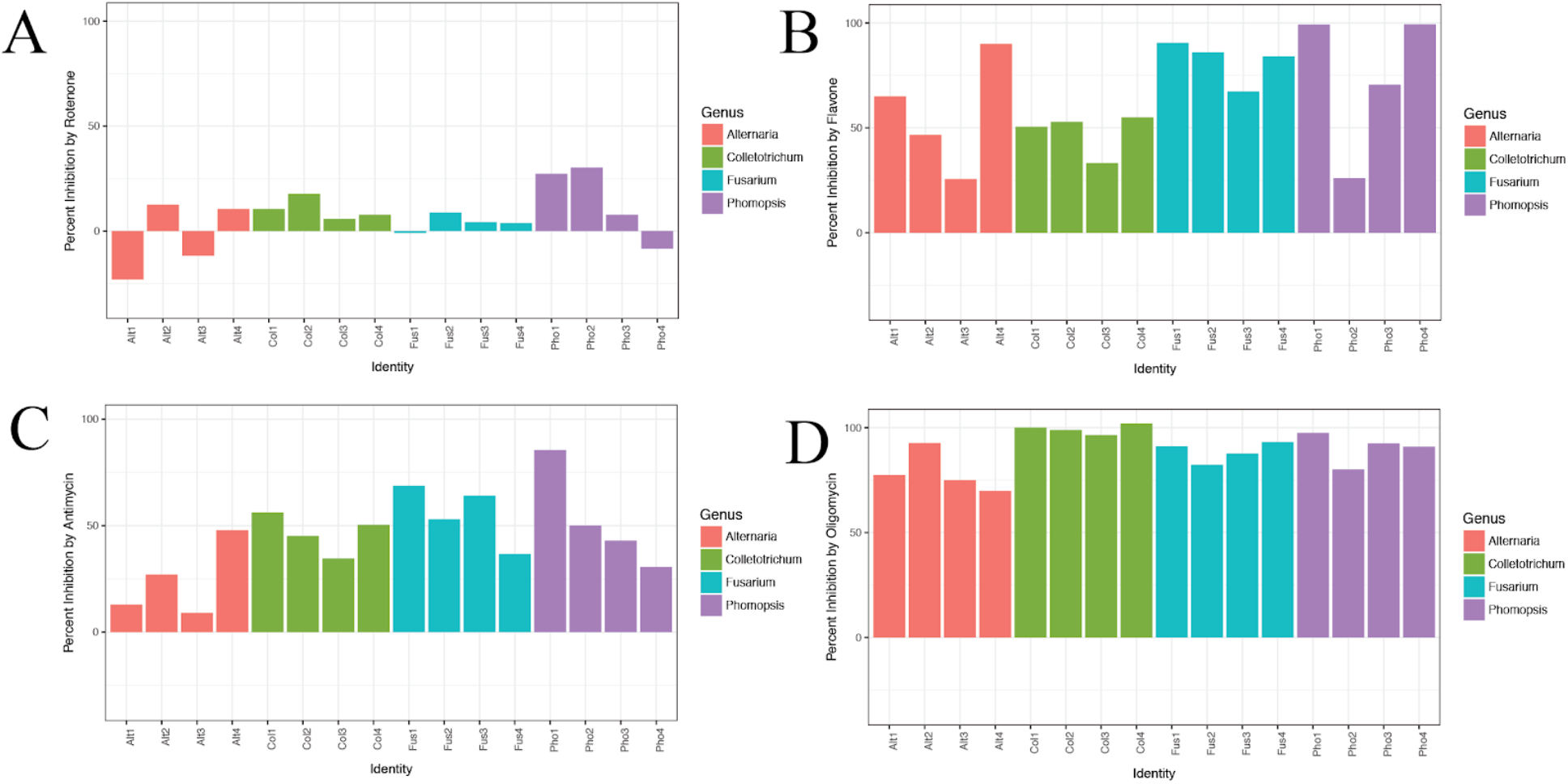
Percent inhibition on glycerol by various drugs. From left to right, red isolates are *Alternaria*, green are *Colletotrichum*, blue are *Fusarium*, and purple are *Phomopsis*. Error bars represent the error propagation standard deviation. **A)** Percent inhibition on glycerol by rotenone, a Complex I inhibitor. Average inhibition across isolates was low at 4.83%, indicating the presence of one or more alternative Complex I enzymes. **B)** Percent inhibition on glycerol by flavone, an inhibitor of multiple targets in the ETC, including alternative NADH dehydrogenases. Average inhibition across isolates was 67.69%. Partial sensitivity confirms the presence of alternative Complex I enzyme(s). **C:** Percent inhibition on glycerol by antimycin, a Complex III inhibitor. Average inhibition across isolates was 44.30%, indicating the presence of an alternative oxidase. **D:** Percent inhibition on glycerol by oligomycin, an ATP synthase inhibitor. Average inhibition across isolates was high at 80.00%, indicating no alternative ATP synthase.

Because inhibition by flavone was low in these chili fungi relative to microbes used in many other studies (Seo et al. 1998), we tested inhibition by Complex II inhibitor antimycin. On glycerol, average percent inhibition by antimycin was 44.30%, indicating partial sensitivity (Figure 2C), implying the presence of an alternative oxidase in many isolates.

To confirm this result, we tested if the inhibition by antimycin correlated with inhibition by flavone, which inhibits alternative Complex I enzymes. Inhibition by antimycin was positively correlated with inhibition by flavone before adjusting for multiple comparisons (p = 0.045), but after adjustment was not (p = 0.085). However, after adjusting for multiple comparisons its effects positively correlated with those of the broad-spectrum ETC inhibitor chloramphenicol (Figure 3) (R^2^ = 0.750, p = 0.0008), supporting the hypothesis that the fungi possess an alternative oxidase. Correlations of all possible drug combinations were also calculated, and their corresponding R^2^ values are shown in Supplementary Table 1.

**Figure 3.**
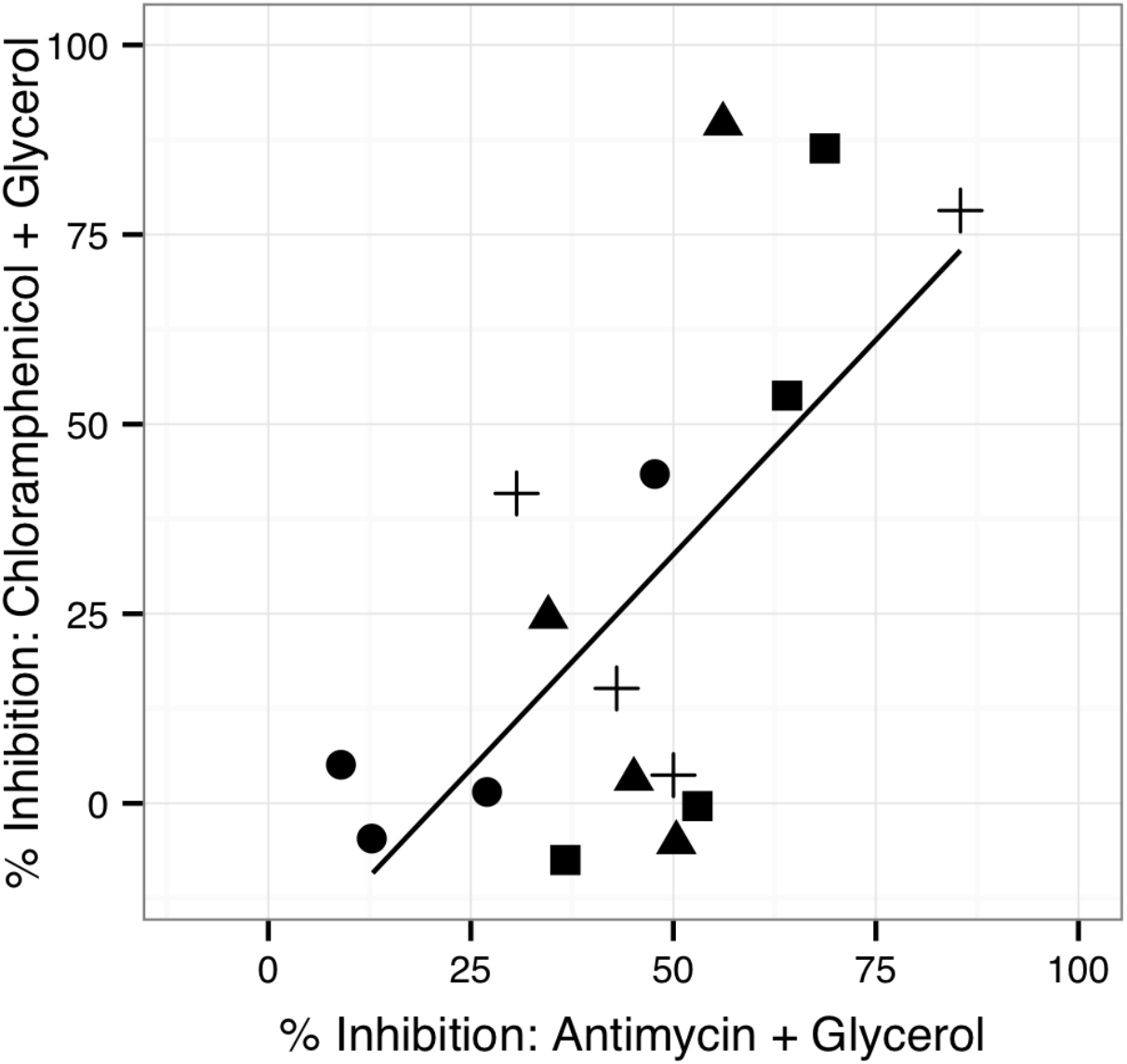
Further evidence of an alternative oxidase. Percent inhibition on glycerol by Complex III inhibitor antimycin is positively correlated with percent inhibition on glycerol by the broad-spectrum ETC inhibitor, chloramphenicol, suggesting the presence of an alternative oxidase. *Alternaria* species are denoted by circles, *Colletotrichum* by triangles, *Fusarium* by squares, and *Phomopsis* by crosses. R^2^ = 0.4254, p = 0.006176.

To rule out the possibility of an alternative ATP synthase, we tested growth in the presence of the ATP synthase inhibitor, oligomycin. This drug was the most effective of the drugs administered, with an average inhibition of 80.00% (Figure 2D), ruling out the presence of an alternative ATP synthase.

In addition to causing the highest overall inhibition, oligomycin was the only drug to cause a significantly higher percent inhibition on fermentable glycerol compared to non-fermentable glucose (Figure 4) (15% higher inhibition on glycerol, one sided t-test, p = 0.03674).

**Figure 4.**
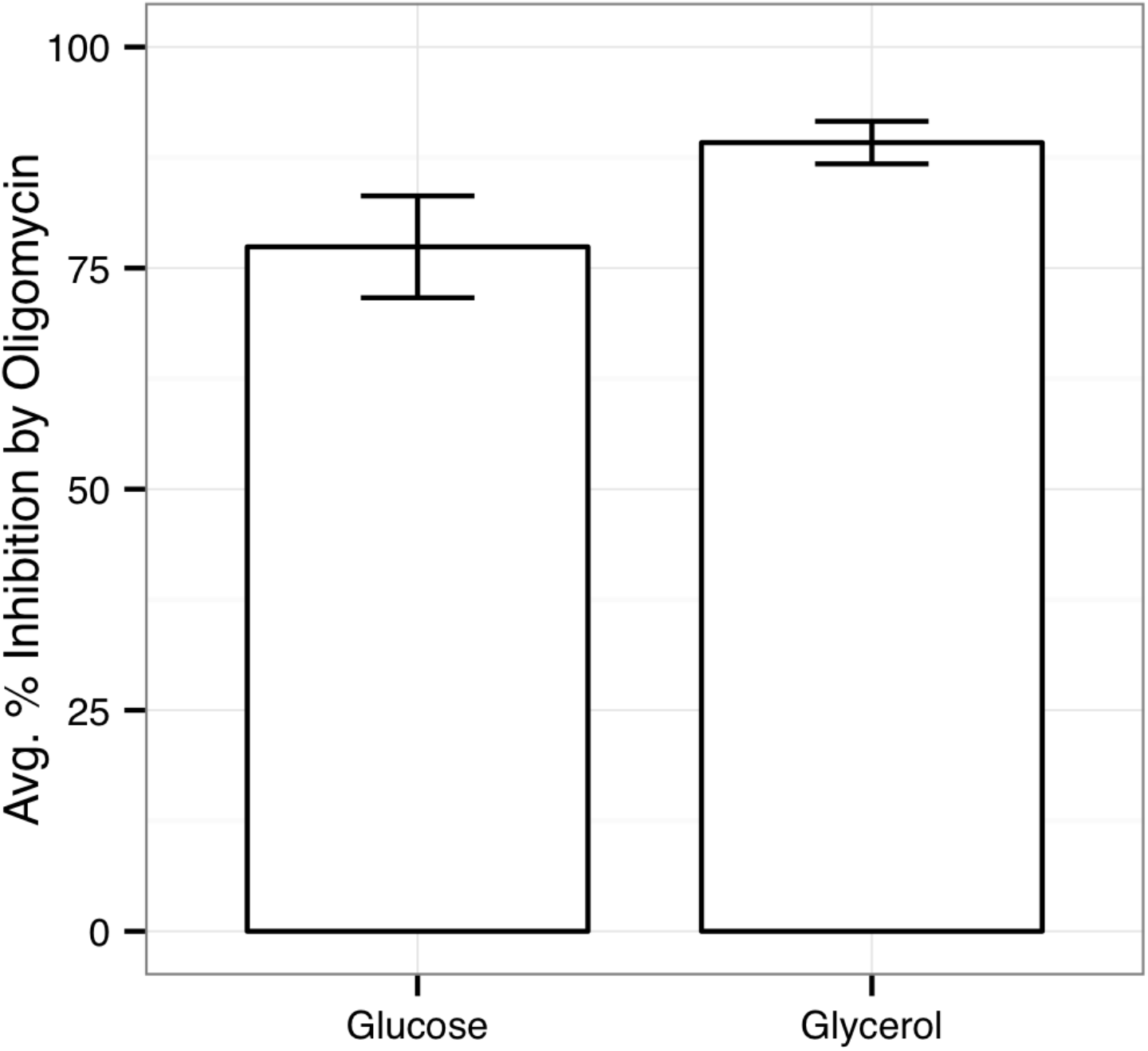
Effects of the ATP synthase inhibitor oligomycin are carbon-dependent. Inhibition by oligomycin was greater on glycerol than glucose, and percent inhibition was 15% higher on glycerol, the carbon source that cannot be fermented (one-sided t test, p = 0.03674. Error bars show standard deviation).

### Presence of alternative respiratory enzymes comes at a fitness cost to basal growth rate

Because the alternative respiratory enzymes are less efficient at energy production compared to the conventional ETC complexes, we next tested if tolerance to any drugs came at a fitness cost to basal growth rate. To do so, we compared percent the inhibition of each drug on both glycerol (non-fermentable) and glucose (fermentable), without drugs. A positive correlation between these two values would indicate that the enzymes used to confer tolerance in the presence of a drug also retard basal growth rate in the absence of drugs.

The only drug that showed a significantly positive correlation was the broad-spectrum ETC inhibitor, flavone (Figure 5) (R^2^ = 0.2826, p = 0.034, values were adjusted for multiple comparisons), indicating that tolerance to flavone comes at a cost to basal growth rate.

**Figure 5.**
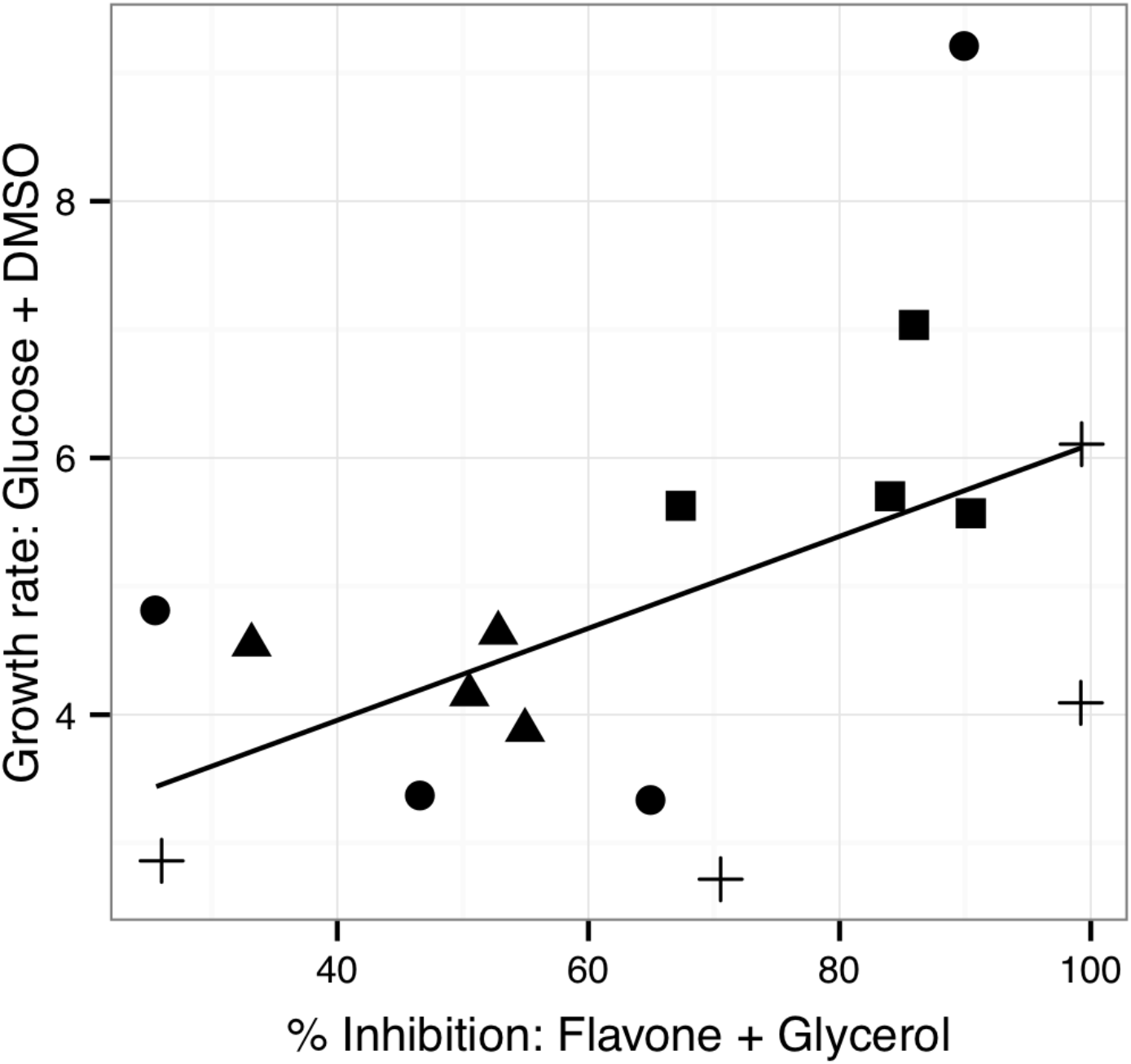
The cost of tolerance to flavone. The lower the inhibitory effects of flavone on an isolate, the slower that isolate is able to grow on glucose in the absence of drugs (controlling for effects of the drug solvent DMSO). *Alternaria* species are denoted by circles, *Colletotrichum* by triangles, *Fusarium* by squares, and *Phomopsis* by crosses (R^2^ = 0.2826, p = 0.03406).

### Fungal pathogens of wild chili peppers are capable of capsaicinoid breakdown

In addition to alternative respiratory enzymes, fungi may possess enzymes that can breakdown capsaicinoids, reducing their concentration. The addition of capsaicinoids to minimal media renders the media cloudy (Supplementary Figure 1A), and precipitates mm-long flavone crystals. These effects allow for the observance of potential zones of clearance on the capsaicin, dihydrocapsaicin, and flavone treatments.

To validate the plate-based assays, we also measured the total concentration of capsaicin in plates with fungal growth via LC-MS. After 10 days of growth, over 99% of the capsaicin had disappeared relative to uninoculated controls (Figure 6). Previous work has reported potential routes of capsaicin catabolism (Price et al. 2019; van den Heuvel et al. 2001), but current LC-TOF analysis of spent media could not detect specific masses consistent with predicted degradation products, such as vanillylamine, vanillin, or vanillate (Supplementary Figure 2A). In Fungal capsaicin plates, a large peak appeared at RT 6.5 that was unique to that condition (Supplementary Figure 2B), and contained a compound with observed m/z of 137.0610. A search of chemicals compatible with this m/z revealed the compound 2-methoxy-4-methylenecyclohexa-2,5-dien-1-one (predicted mass 137.0597), which may represent a novel metabolite formed in capsaicin degradation.

**Figure 6.**
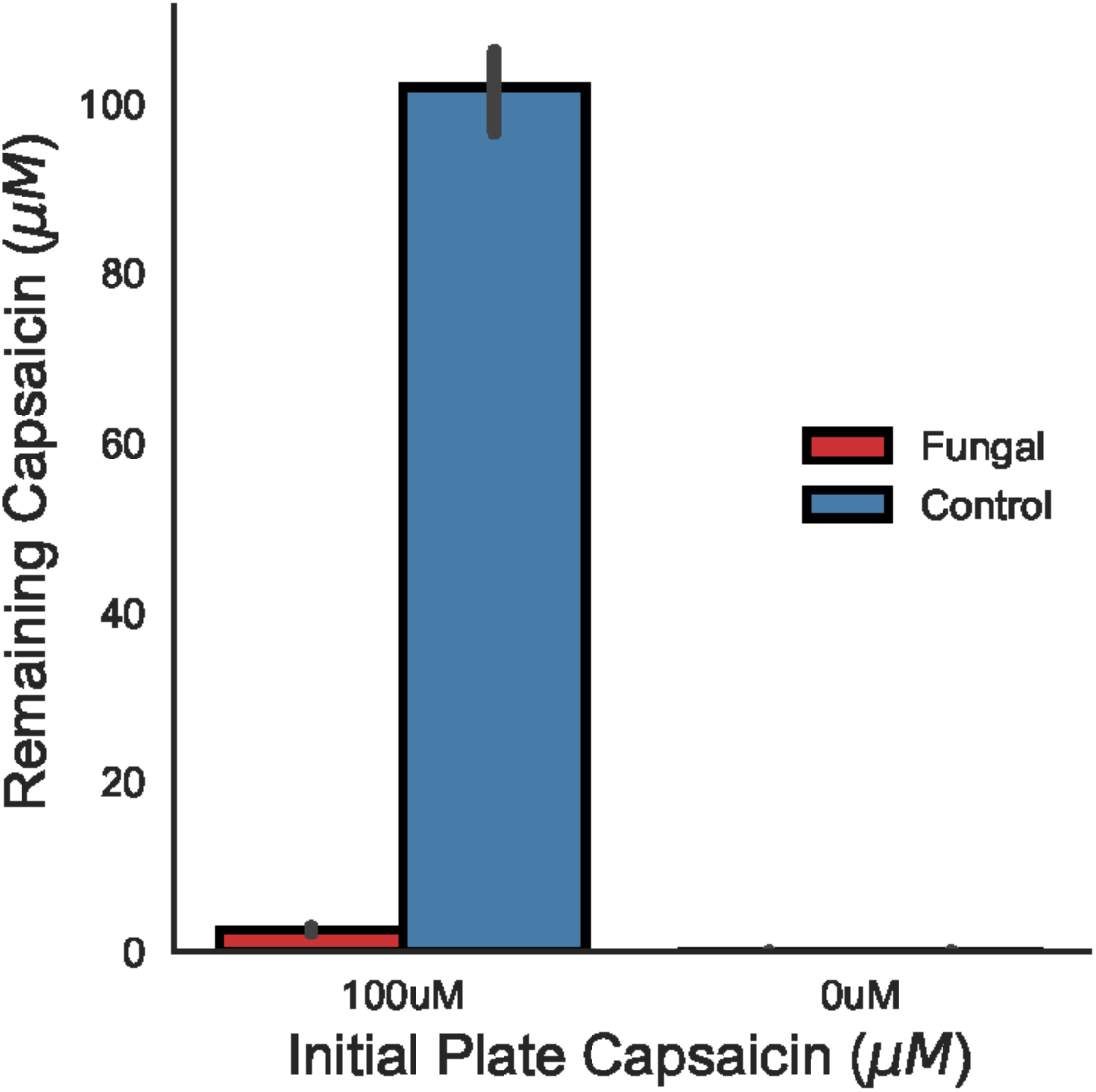
LC-MS analysis of capsaicin degradation. LC-MS analysis confirmed the capsaicin was degraded in Fungal plates, but not Controls (Two-sided t-test, p ⩽ 0.001, n = 3. Error bars are 95% Confidence Intervals).

### OXPHOS inhibition alone does not explain the inhibitory effects of capsaicinoids

Lastly, we tested if the capsaicinoids’ inhibition of components of the electron transport chain explained the overall inhibitory effects of either drug. To do this, we first determined how the fungi primarily obtain energy. Because glycerol can only be used in OXPHOS and glucose can be used in either OXPHOS or fermentation, a positive correlation between growth rate on the two carbon sources would indicate that, given the choice between OXPHOS and fermentation, the fungi primarily use OXPHOS.

To examine the roles of the capsaicinoids on the ETC, we compared the percent inhibition by each drug on glycerol compared to glucose. Because these fungi generate a majority of their energy via OXPHOS (Figure 7), a positive correlation between percent inhibition by a drug on glycerol and percent inhibition on glucose indicates the drug inhibits OXPHOS, but does not noticeably affect other aspects of metabolism. A lack of correlation indicates the drug has significant effects on more components than the electron transport chain, or that they are unable to ferment.

**Figure 7.**
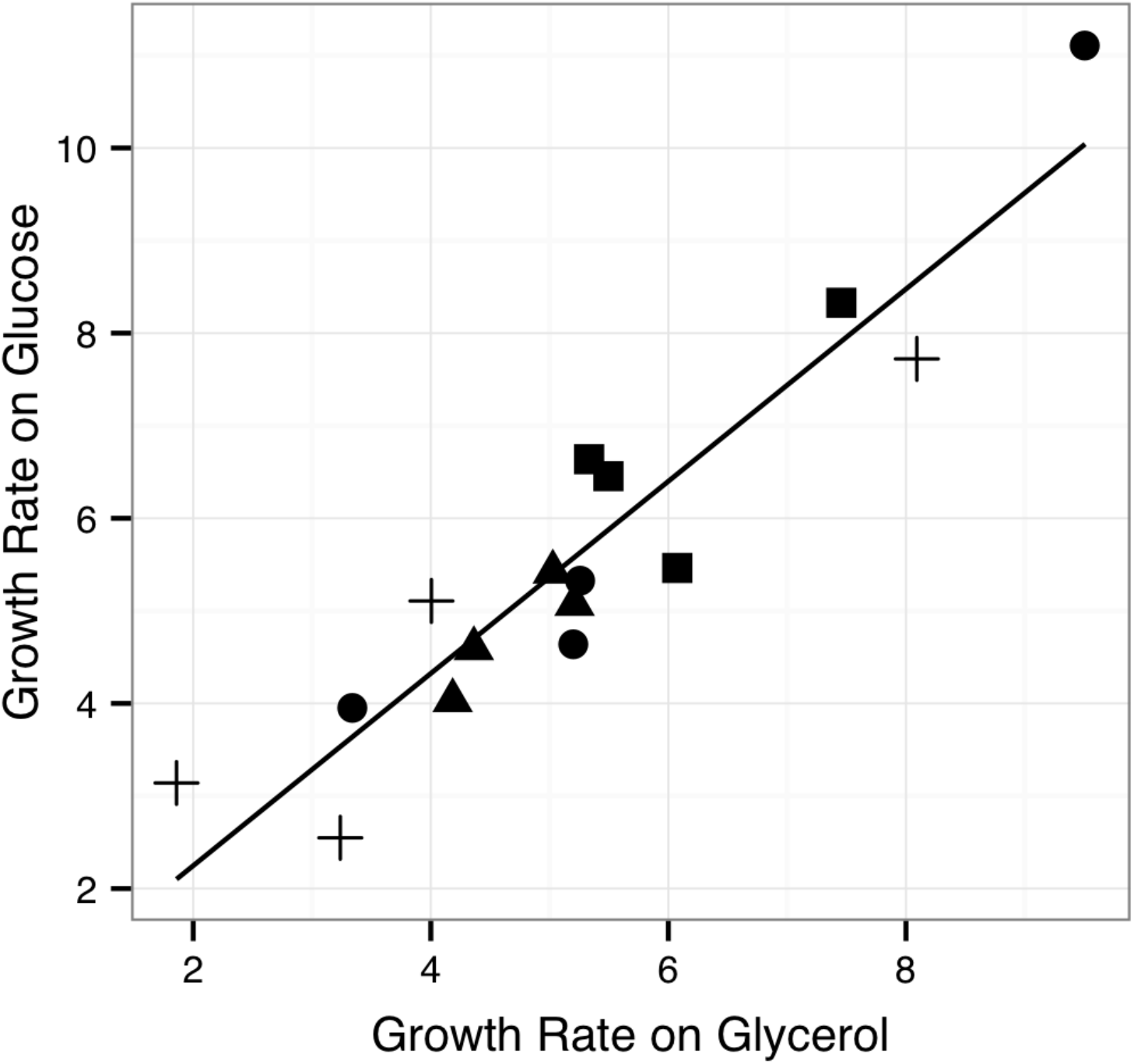
Fungal growth rate in the absence of drugs. Growth rate on glucose strongly correlates with growth rate on glycerol. Under normal conditions, these fungi use oxidative phosphorylation to produce the majority of their energy. *Alternaria* in circles; *Colletotrichum* in triangles; *Fusarium* in squares; *Phomopsis* in crosses (R^2^ = 0.8736, p ⩽ 0.001).

We found a weakly positive correlation between percent inhibition on glucose and glycerol for dihydrocapsaicin (Figure 8A) (R^2^ = 0.455, p = 0.0225), but did not find a significant correlation between percent inhibition on the different carbon sources for capsaicin (Figure 8B) (R^2^ = 0.01105, p = 0.2981). From this we conclude dihydrocapsaicin likely solely affects the ETC, while capsaicin exerts additional effects on the cell.

**Figure 8.**
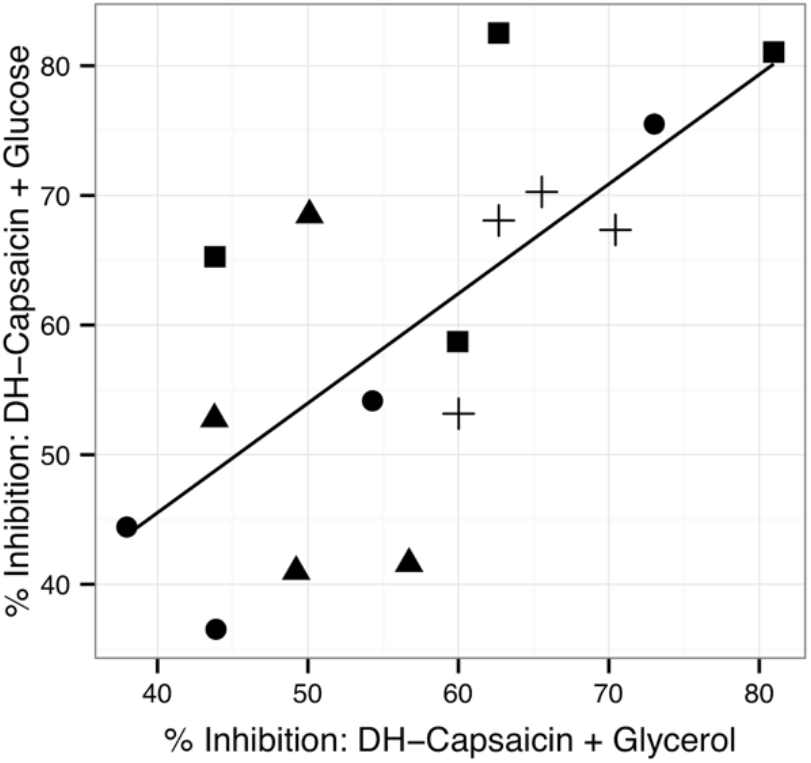
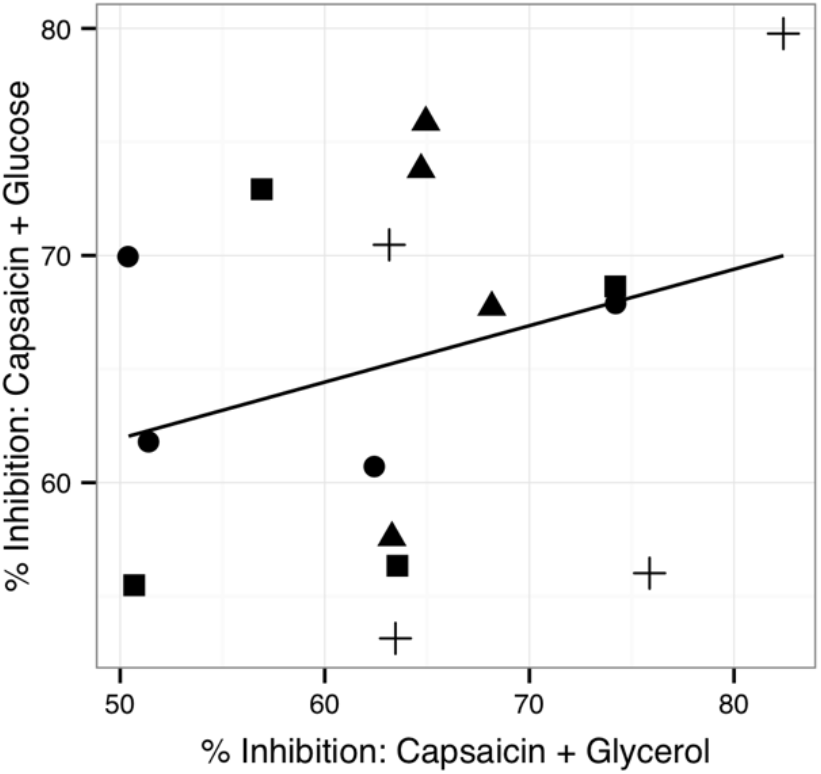
Capsaicinoids do not exert effects outside of the ETC. A) OXPHOS inhibition alone explains the inhibitory effects of dihydrocapsaicin (R^2^ = 0.45, p = .0025). B) OXPHOS inhibition alone does not explain the inhibitory effects of capsaicin (R^2^ = 0.07698, p = 0.2981).

## Discussion

Here we found fungal pathogens of wild chili peppers, which have evolved in the presence of capsaicinoids, possess multiple means to tolerate capsaicinoids. Firstly, all fungal isolates tested were insensitive to the Complex I inhibitor rotenone, indicating all isolates possess at least one alternative NADH dehydrogenase (Figure 2A)(Joseph-Horne et al. 2001; Juárez et al. 2004; Kerscher et al. 1999).The fungal species used include two fungal genera in which alternative Complex I enzymes had been previously inferred from published genomic data, *Fusarium* and *Colletotrichum*, in addition to two genera not previously described as possessing respiratory alternatives, *Phomopsis* and *Alternaria*.

The phylogenetic breadth of these alternative Complex I enzymes in these fungi suggests alternative NADH dehydrogenases may be more widespread in fungal phytopathogens than previously believed. Plant fruits are often rich in secondary metabolites that target the electron transport chain. In *Aspergillus niger*, the alternative NADH dehydrogenase is less efficient at energy production than the classical enzyme, but serves in an additional protective role against oxidative stress. Recognition of fungal alternative NADH dehydrogenases has already resulted in the creation of novel drugs that specifically target them (Mogi et al. 2009). Alternative NADH enzymes are not present in mammals, making them an excellent target for treating opportunistic fungal infections in humans, pets, and livestock.

In other genetically intractable fungi, the presence of an alternative NADH has been inferred by the lack of sensitivity to rotenone coupled with the presence of sensitivity to flavone (Tudella et al. 2004; Uyemura et al. 2004). Though the fungal isolates tested here were highly tolerant to rotenone (Figure 2A), they showed an unanticipated range of sensitivities to flavone (Figure 2B). The range of sensitivities to the Complex III inhibitor antimycin (Figure 2C), combined with the tolerance to flavone, suggests many isolates (e.g. Alt1, Alt 3, Col3, Fus 4, and Pho4) possess an alternative cytochrome oxidase as well (Descheneau et al. 2005; Affourtit et al. 2000).

The effects of other inhibitors bolster support for the presence of an alternative oxidase in some isolates. The inhibition of antimycin correlated significantly with inhibition by chloramphenicol, a broad-spectrum ETC inhibitor that targets multiple standard and alternative enzymes such as the alternative oxidase, and decreases total mitochondrial translation (Table 1; Figure 3, R^2^ = 0.43, p = 0.006) (Descheneau et al. 2005). In particular, chloramphenicol disrupts translation of mitochondrial complex III. As Descheneau et al observe, previous researchers have misinterpreted the importance of chloramphenicol resistance as general drug resistance, when the reason for the resistance is the presence of a specific alternative oxidase. When genome data of a microbe of interest is not available, drug tolerance can be a useful screen for the presence of alternative respiratory enzymes.

Alternative oxidases have been described in the filamentous fungi *Neurospora crassa (Descheneau et al. 2005)*, *Aspergillus fumigatus* (Tudella et al. 2004), *Fusarium oxysporom (Pereira et al. 1997)* and others. In fungi such as *Ustilago maydis*, the alternative oxidase is not necessary for normal growth, but confers a fitness advantage in the presence of respiratory stress (Cárdenas-Monroy et al. 2017). Fungi that associate with spicy chili pepper fruits experience a substantial degree of respiratory stress induced by capsaicinoids, but that stress may be alleviated by an alternative oxidase.

We found evidence for alternative oxidases in three additional genera: *Alternaria, Colletotrichum*, and *Phomopsis*. Much like alternative NADH dehydrogenases, alternative oxidases may be more prevalent among wild plant pathogens than previously believed. However, while this work is strong in its ability to detect ecologically relevant patterns, it is weak in terms of molecular precision. Much work remains to isolate the proteins inferred here, and to perform knock-out studies to test how much each enzyme contributes to capsaicinoid tolerance. Further work with the fungal isolates will aim to elucidate the structure and function of these enzymes.

Here, the variation in flavone inhibition (Figure 2B) may be partially due to flavone’s action on other enzymes in the cell. In humans, flavonoids bind to a broad range of enzymes including human topoisomerase I (Boege et al. 1996), cyclin-dependent kinases (CDKs) (De Azevedo et al. 1996), phosphatidylinositol 3-kinase alpha (Agullo et al. 1997), and xanthine oxidase (Cos et al. 1998). In isolated rat liver mitochondria, the flavonoids quercetin and galangin interacted with the mitochondrial membrane (Dorta et al. 2005). Dorta et al attribute some of the compounds’ activity to the double bond between the ether and ketone of the middle C-ring, which is also present in flavone. Flavone’s ability to affect a variety of cell components could help explain why inhibition by antimycin did not correlate with inhibition by flavone, but did correlate with that of chloramphenicol; the effects of chloramphenicol are more targeted to mitochondrial protein synthesis (Juárez et al. 2004) and the ETC (Zhang et al. 1999; Vinnikov et al. 1994).

Tolerance to drugs can come at a cost to basal growth rate. Here we observed such a tradeoff when examining tolerance to flavone (Figure 5): the lower the inhibitory effects of flavone on an isolate, the slower that isolate grew on glucose in the absence of ETC-inhibiting drugs. Flavone binds to multiple ETC complexes, and can bind to both the external (Juárez et al. 2004) and internal NADH enzymes (Seo et al. 1998). To use OXPHOS in the presence of flavone, fungi must engage both the alternative NADH dehydrogenase(s) and the alternative oxidase, which transfer electrons down the ETC, but do not pump protons into the inter membrane space of the mitochondria, lessening the proton motive force (Joseph-Horne et al. 2001). For fungi that associate with plants which produce high titers of electron transport chain disruptors, such as chili peppers, the ETC disruptors may exact a higher selective pressure than basal growth rate. More research interest in the prevalence and action of alternative respiratory enzymes in fungi will inform our understanding of their evolutionary and ecological significance. This knowledge can be leveraged in direct applications such as plant pathology and food crop contamination, and also to aid medical efforts to treat opportunistic fungal infections (Martins et al. 2011).

Oligomycin, the ATP synthase inhibitor, had the highest percent inhibition of any drug, and the lowest standard deviation (Figure 2D). This is likely because, of all the ETC complexes, the ATP synthase is the largest: it is a supercomplex weighing in at over 500kDa, composed of two separate rotary motors (Okuno et al. 2011). Because the ATPase consists of so many separate proteins, the synthase would be especially difficult for natural selection to duplicate. In a recent analysis of fungal genomes, there were no duplications of the ATP synthase genes (Marcet-Houben et al. 2009). Targeting the ATP synthase may be an effective strategy for slowing fungal pathogens that possess multiple alternative enzymes.

Furthermore, oligomycin caused 15% more inhibition on the carbon source glycerol, which can not be fermented, compared with the fermentable sugar, glucose (Figure 4). We propose drugs that strongly inhibit the electron transport chain in phytopathogenic fungi can cause different effect sizes depending on the carbon source used. This effect was found in human fibroblasts (Gohil et al. 2010), but has not, to our knowledge, been shown in fungal plant pathogens. We advocate for mindful selection of carbon sources when testing the efficacy of drugs on plant pathogens *in vitro*. Moreover, knowledge of which mode of energy production fungi favor can inform applied research with plant pathogens *in vivo* as well. For example, the observation that cancerous cells prefer to ferment (Warburg 1956) has led to novel treatments that target glycolysis in tumor cells (Diaz-Ruiz et al. 2011). In agriculture, applying combinations of fungicides to target both fermentation and OXPHOS may be an effective strategy to limit crop loss and improve yield.

The present results also show tolerance to capsaicin may involve mechanisms outside the ETC. Further resistance to capsaicinoids may be due to these fungal isolates’ ability to degrade capsaicinoids, as was inferred by a clear zone around many of the fungal plugs (Supplementary Figure 1A) and confirmed by LCMS (Figure 6). While the degradation of capsaicinoids has been studied in other organisms, in these previous studies the capsaicinoids are always broken into vanillylamine, which we did not specifically identify (Supplementary Figure 2A). Here, the fungi are either immediately modifying vanillylamine into the detected products (Figure S2B, or they are breaking down capsaicinoids using a novel, hitherto undescribed pathway. Future work will explore the structure and function of the enzymes involved in this pathway, and possible applications such as treating accidental applications of mace or de-spicing food.

We designed an assay to test whether we could rule out the small number of studies purporting that capsaicin disrupts cell membranes. After confirming that our isolates generate the majority of their energy via OXPHOS (Figure 7), we tested the inhibition of the two primary capsaicinoids on the two carbon sources. We found the effect of capsaicin on OXPHOS did not explain total inhibition of capsaicin (Figure 8B). Effect on OXPHOS, however, did weakly explain inhibition by dihydrocapsaicin, the second most common capsaicinoid in chili peppers (Figure 8A). The two compounds are each roughly the length of half a phospholipid bilayer, and the only structural difference between them is that capsaicin has a double bond in the hydrophobic tail (Supplementary Figure 2B, 2C). This nonrotating double bond makes capsaicin more rigid than dihydrocapsaicin, which, in a cell, might cause more membrane disruption and subsequent cell content leakage.

Determining the potential effect of capsaicin on cell membranes is important not only for quantifying the total effect of capsaicin on microbes, but also for human medicine. Capsaicin is used in a number of medical applications, ranging from topical creams to treat human ailments such as diabetic neuropathy, psoriasis, and rheumatoid arthritis (Reyes-Escogido et al. 2011), to treatment of various types of cancer such as skin cancers, colon cancer, and lung cancer (Bley et al. 2012). At the same time, research has hinted that capsaicin itself can be a carcinogen (López-Carrillo et al. 2003), and researchers have yet to reach a conclusion on whether this compound is more often beneficial or harmful to human health (Bode and Dong 2011). While the binding of capsaicin to human TRPV receptors has been studied in detail (Szallasi et al. 2007), the ability of capsaicin to disrupt human cell and mitochondrial membranes has been largely ignored, and demands immediate research.

In summary, our data suggest fungal seed pathogens of wild chili peppers possess alternative Complex I enzymes that allow them to produce energy in the presence of capsaicinoids. We also found evidence for an alternative oxidase, suggesting plant pathogens may possess more alternative respiratory enzymes than were previously known, and may have evolved these enzymes in response to secondary metabolite production by plants. Fungal growth inhibition by dihydrocapsaicin is explained through dihydrocapsaicin’s effects on OXPHOS, but OXPHOS inhibition alone does not explain the total effects of capsaicin. Our results suggest the separate bodies of work regarding capsaicin’s effects on microbes are actually complementary: capsaicin likely affects both respiratory enzymes *and* cell membranes. To tolerate capsaicin, fungi employ a variety of strategies, including multiple alternative respiratory enzymes and capsaicin degradation. This work offers practical applications for the protection of *Capsicum* and other crops against seed pathogens, presents implications for human medicine, and contributes to the evolutionary and ecological theories of plant-pathogen evolution.

## Acknowledgments

We are indebted to Tristan Wang for his detailed measuring at the microscope, Vamsi Mootha for help with experimental design, Steve Worthington for statistical advice, and to the members of the Pringle and Bruns/Taylor Labs for help with manuscript revisions. C Adams would like to thank C. O. Bodom for inspiration. The gifts of the culture collection, capsaicin, and dihydrocapsacin from Joshua J Tewksbury are gratefully acknowledged.

This work was funded in part by Harvard University, the University of California-Berkeley, the National Science Foundation Graduate Research Fellowship, and the University of California-Berkeley Chancellor’s Fellowship.

---

**Supplementary Figure 1.**
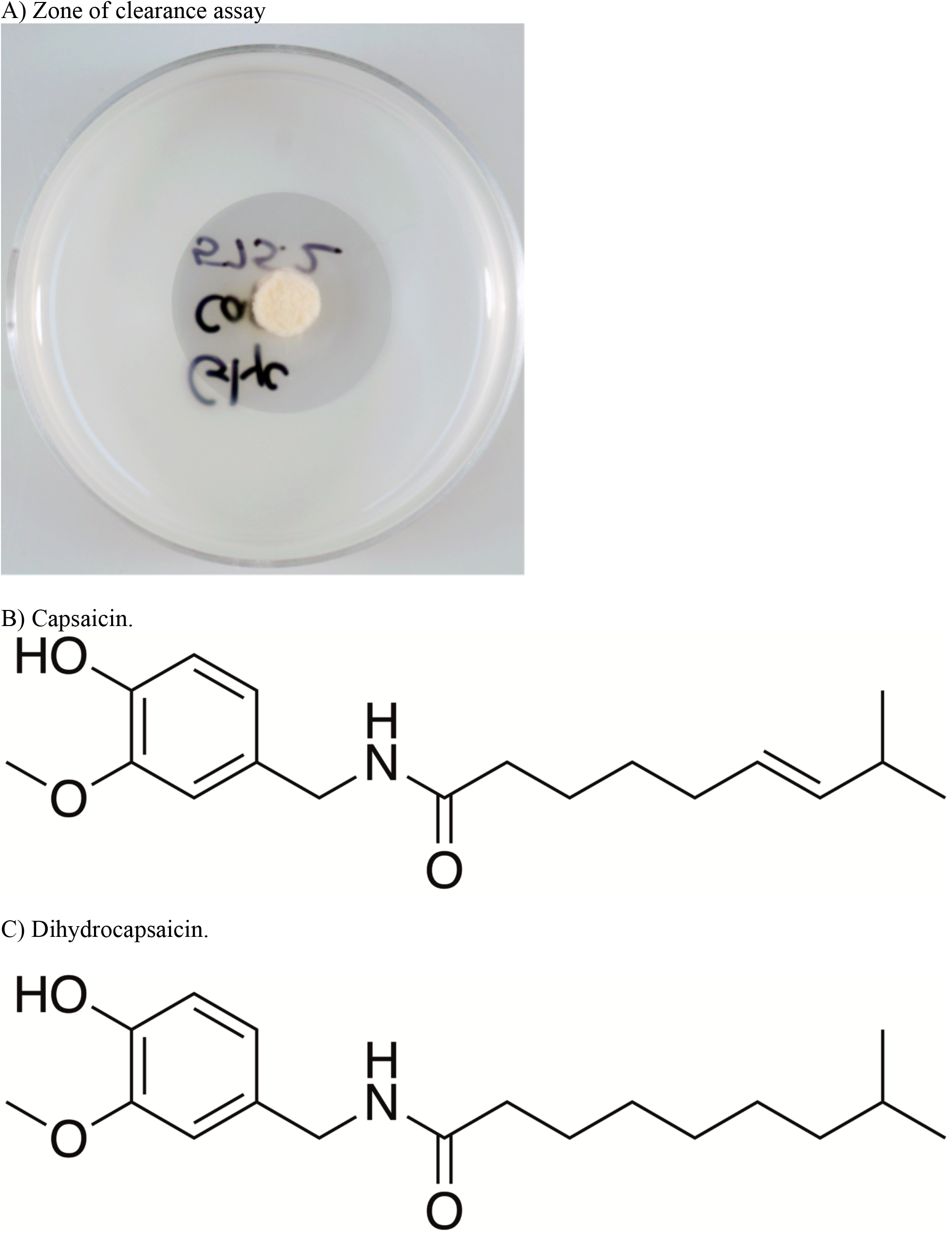
Zone of clearance and compound structures. A) The zone of clearance outside the fungal growth area. Clearance zones were observed when isolates were grown on capsaicin, dihydrocapsaicin, and flavone. B) the structure of capsaicin and C) dihydrocapsaicin. The two compounds are structurally identical except for the double bond at the 6th carbon of the hydrophobic tail, rendering capsaicin more rigid than dihydrocapsaicin, and more likely to disrupt membranes.

**Supplementary Table 1.**
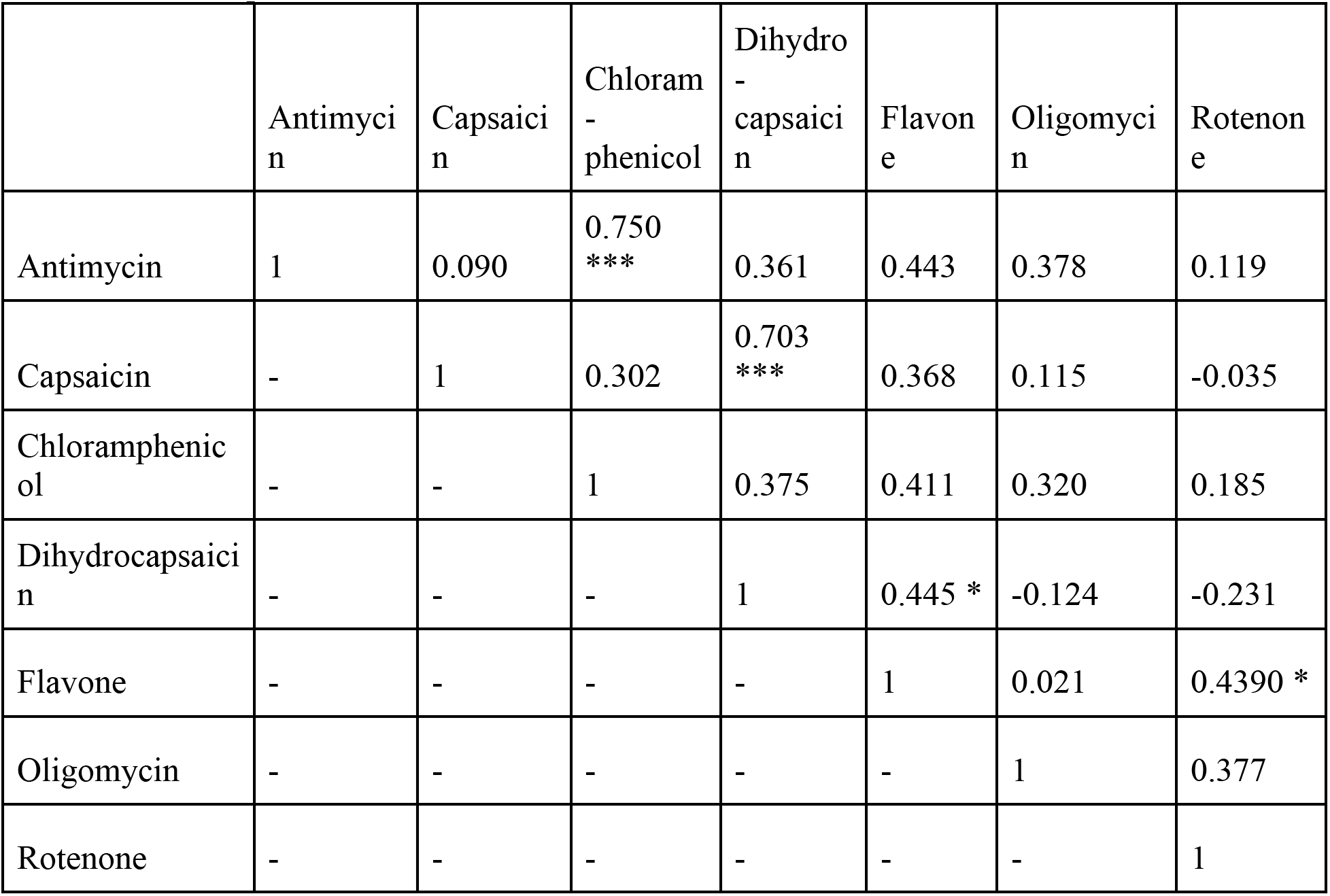
Results of analysis of correlations of percent inhibition on glycerol by all drug combinations. Statistically significant correlations (p < .05) are marked with ***. Correlations with p < .1 are marked with *.

|  | Antimycin | Capsaicin | Chloramphenicol | Dihydrocapsaicin | Flavone | Oligomycin | Rotenone |
| --- | --- | --- | --- | --- | --- | --- | --- |
| Antimycin | 1 | 0.090 | 0.750 *** | 0.361 | 0.443 | 0.378 | 0.119 |
| Capsaicin | - | 1 | 0.302 | 0.703 *** | 0.368 | 0.115 | -0.035 |
| Chloramphenicol | - | - | 1 | 0.375 | 0.411 | 0.320 | 0.185 |
| Dihydrocapsaicin | - | - | - | 1 | 0.445 * | -0.124 | -0.231 |
| Flavone | - | - | - | - | 1 | 0.021 | 0.4390 * |
| Oligomycin | - | - | - | - | - | 1 | 0.377 |
| Rotenone | - | - | - | - | - | - | 1 |

